# On the inconsistency of pollinator species traits for predicting either response to agricultural intensification or functional contribution

**DOI:** 10.1101/072132

**Authors:** Ignasi Bartomeus, Daniel P. Cariveau, Tina Harrison, Rachael Winfree

## Abstract

The response and effect trait framework, if supported empirically, would provide for powerful and general predictions about how biodiversity loss will lead to loss in ecosystem function. This framework proposes that species traits will explain how different species respond to disturbance (i.e. response traits) as well as their contribution to ecosystem function (i.e. effect traits). However, predictive response and effect traits remain elusive for most systems. Here, we present detailed data on crop pollination services provided by native, wild bees to explore the role of six commonly used species traits in determining how crop pollination is affected by increasing agricultural intensification. Analyses were conducted in parallel for three crop systems (watermelon, cranberry, and blueberry) located within the same geographical region (mid-Atlantic USA). Bee species traits did not strongly predict species’ response to agricultural intensification, and the few traits that were weakly predictive were not consistent across crops. Similarly, no trait predicted species’ overall functional contribution in any of the three crop systems, although body size was a good predictor of per capita efficiency in two systems. So far, most studies looking for response or effect traits in pollination systems have found weak and often contradicting links. Overall we were unable to make generalizable predictions regarding species responses to land-use change and its effect on the delivery of ecosystem services. Pollinator traits may be useful for understanding ecological processes in some systems, but thus far the promise of traits-based ecology has yet to be fulfilled for pollination ecology.

## Introduction

Land-use change, along with other human-induced global change drivers, is accelerating the rates of extinction of most taxa (Ellis et al. 2010). At the same time, humanity relies on ecosystem services that wild species deliver, such as pollination and pest control by insects, and nutrient cycling by microorganisms (Cardinale et al. 2012). Thus, it is important to understand the relationship between biodiversity loss and ecosystem service delivery (Schwartz et al. 2000). In particular, making generalizable predictions regarding how the decline or local extinction of taxa will affect ecosystem services will allow for targeted conservation actions to ameliorate negative impacts of land-use change.

One avenue for predicting the functional consequences of biodiversity loss is the response and effect trait framework (Lavorel and Garnier 2002, Naeem and Wright 2003, McGill et al. 2006). Local extinction does not occur at random because extinction risk is dependent on the species’ characteristics. Identifying which traits govern species responses to particular threats (‘response traits’) would provide the first step for predicting future species loss. Furthermore, the magnitude by which ecosystem function declines when a species is lost depends on that species’ functional contribution. This, too, is likely to be mediated by the species’ traits (‘effect traits’). Therefore, the relationship between response and effect traits will mediate the magnitude of the impact of human disturbance on ecosystem services (Schleuning et al. 2015). For example, if the same species traits that are associated with high function are also most sensitive to disturbance, ecosystem function would be predicted to decline rapidly (Larsen et al. 2005).

However, for the response-effect trait framework to be useful, it is first necessary to identify response and effect traits that are both explanatory and possible to measure in the field (Cadotte et al. 2011). While a few generalities have emerged as to which traits make animal species at greater risk of local decline, including dietary or habitat specialization and body size (Fisher and Owens 2004, Flynn et al. 2009, Öckinger et al. 2010), the correlation between these response traits and extinction risk has been found to be weak, variable, or context-dependent (Devictor et al. 2008, Fritz et al. 2009, Powney et al. 2014). Similarly, although some effect traits have been identified, they are often weakly predictive, and their identity varies by function and taxonomic group (Gagic et al. 2015). Lastly, within the functional trait field as a whole, most progress has been made in identifying functional traits for plants (Díaz et al. 2016), while little is known for animals (Didham et al. 2016).

Here, we seek to identify response and effect traits for wild bee species providing a key ecosystem service, crop pollination. The yield of most crop plants increases with animal pollination (Klein et al. 2007). While managed honey bees are a leading crop pollinator, wild insects contribute more than half of pollinator visits to crop flowers across more than 40 crop systems worldwide (Rader et al. 2016). A major threat to pollinators is habitat destruction, primarily conversion to agriculture (Garibaldi et al. 2011), which is also a leading cause of species loss worldwide (Pereira et al. 2010). Thus agricultural land use has the potential to affect one of the ecosystem services upon which agriculture itself depends (Deguines et al. 2014).

Our data sets were collected and analyzed in parallel and come from three crop systems (watermelon, cranberry and blueberry) located within the same geographical region (mid-Atlantic USA), but pollinated by distinct bee communities. We determined whether six commonly-used species traits can predict 1) species’ responses to agricultural intensification (response traits) and/or 2) species’ contributions to crop pollination (effect traits) and discuss our results on the light of recently published studies on pollinator environment–trait and pollination-trait associations.

## Material and methods

### Study system

We selected 49 sites across three study systems that were located throughout New Jersey and eastern Pennsylvania (USA). Watermelon sites (N = 17) were located in 90 × 60 km region central New Jersey and Eastern Pennsylvania, where the main types of land use are agriculture and suburban development, interspersed with highly fragmented deciduous forest. Cranberry and blueberry sites (N = 16 each) were both located within a 35 x 55 km area in southern New Jersey, where the main land cover types are pine-oak ericaceous heath and agriculture. All sites in all systems were separated by at least 1 km (range, watermelon: 2-90 km, cranberry: 1-32 km, blueberry 1-38 km).

All three crops are highly dependent upon bee pollination for marketable fruit production (Klein et al. 2007). Commercial honey bees are used in most of our study fields. However, honey bees are primarily managed hives, moved throughout the region, and only found on sites during bloom. Therefore, honey bees are not influenced by land cover in the same manner as wild bees and are not used in our analyses. Wild bees are important pollinators in all three systems (mean percentage of wild bee visits: 73% watermelon, 25% cranberry, and 14% blueberry).

### Data collection

At all sites on all three crops, we used hand-netting to measure overall bee abundance and species richness. To collect bees, we walked along a fixed 50-200 m^2^ transect at standard times of day and collected all bees observed to be visiting flowers. In watermelon and blueberry, bees were netted three times throughout the day for 20 minutes per transect (60 minutes per date per site) and twice each day in cranberry for 30 minutes per transect (120 minutes per date per site). Data were collected during the peak bloom in 2010 (watermelon: July, cranberry: late-May-early July, blueberry: April-early May). Data were collected on three days per site for watermelon and blueberry and two days per site for cranberry. Detailed methods can be found in Cariveau et al. (2013), Benjamin et al. (2014) and Winfree et al. (2015).

### Land cover characteristics of sites

To relate pollinator response traits to agricultural intensification we used a commonly used land-use variable, percent land cover in agricultural production surrounding sites (Fahrig 2013). For this end, we required high-quality land cover data for each pollinator collection site. For the cranberry and blueberry sites in New Jersey, we used a continuous polygon layer classified by visual photograph interpretation into 60 categories, at a minimum mapping unit of 4047 m^2^ (1 acre; GIS Data provided by the New Jersey Department of Environmental Protection). For watermelon sites that extend from central New Jersey into Pennsylvania, we created a similar land cover data layer by manually digitizing Google Earth imagery and visually classifying 15 categories, at a minimum mapping unit of 5,000 m^2^ (1.24 acres). As each crop was analyzed separately, our results are robust to using different land cover data. However, to simplify the interpretation of results for the three crops, we reclassified all land cover data into the following 7 broad categories: agriculture, open managed (for example, mowed grass), open natural or semi-natural (for example, old fields), semi-urban (<30% impervious surface), urban (>30% impervious surface), wooded, and open water.

For each data collection site, we calculated two land cover variables: percent agriculture and percent natural and semi-natural open habitat. We used agricultural land cover as our primary land-use change variable as it is the dominant anthropogenic habitat type in all three study systems (Supplementary Table A1). In addition, we also measured percent of open natural/semi-natural habitat, which although it accounts for only a small proportion of the total land cover (Supplementary Table A1), might be disproportionately important as forage and nesting habitat for bees (Kleijn et al. 2006). We calculated values for this two land cover variables at both a small scale (300 m radius) and a large scale (1500 m radius), which correspond to typical flight distances of small- and large-bodied bees, respectively (Greenleaf et al. 2007).

### Pollinator function

To estimate the pollination services provided per bee species, we measured two variables in the field, flower visitation frequency and per visit efficiency. As variation in visitation frequency may be a function of land use at individual farms, we use species abundances for each species at the site with its highest abundance for each crop. Hence, we assess visitation frequency at its maximum, which represents the optimal visitation frequency for each species.

To measure the pollination efficiency, we quantified single-visit pollen deposition by presenting virgin flowers to individual bees foraging on the target crop. After visitation, we counted the number of pollen grains deposited per flower visit (watermelon) or the number of pollen tetrads with pollen tubes per flower visit (cranberry and blueberry). Because species identification in the field is not possible for most bees and net collecting immediately after visits is generally not possible, for the measurement of pollination efficiency we grouped bees in species groups. Each group consisted of between one and 27 species, with the median number of species per group being 4 species (Supplementary Table A2). Control flowers were left bagged until the end of the field day, and contained few pollen grains (watermelon mean = 3 grains, N = 40 stigmas; cranberry mean = 0 tetrads, N = 82 stigmas; blueberry mean = 2 tetrads, N= 734 stigmas). We used mean number of pollen grains deposited by a single visit group and assigned that value to each of the species in the single visit group. For detailed methods see Cariveau et al. (2013), Benjamin et al. (2014), Winfree et al. (2015).

### Species traits

Bee species vary in a number of traits that are associated with their response to land-use change (Williams et al. 2010). Moreover, these traits will likely affect the pollinator contribution to function, either by modifying its abundance or because they are related to its per capita effectiveness. We obtained detailed natural history data on 6 traits for the 90 bee species in our study: a) sociality (solitary, facultative social, eusocial), b) nesting placement (hole, cavity, stem, wood, ground), c) brood parasite (yes, no), d) body size, e) diet breadth (level of specialization) and f) tongue length.

We obtained the trait data as follows. Species sociality level, nesting behavior and brood parasite status were extracted from the literature (Bartomeus et al. 2013a). Body size (estimated from intertegular span, IT; Cane 1987) was measured in the lab using collected specimens that had been identified to the species level by professional taxonomists. Multiple specimens were measured per species (mean = 6.6 specimens ± 3 S.E.) and the mean across the measured specimens was used as the value for the species. Bee body size also correlates strongly with foraging distance (Greenleaf et al. 2007), and thus is ecologically related to mobility. Tongue length was measured in the lab for 7.7 ± 1.2 SE specimens per species, and the mean across the measured specimens is used. For the 40 specimens for which we cannot obtain a tongue measure, we estimated tongue length from the species’ body size and phylogeny using an allometric equation (Cariveau et al. 2016).

Diet breadth was calculated using six independent datasets previously collected at 139 sites throughout the study region by the Winfree laboratory group. Each data set consists of individual pollinator specimens that were net-collected while foraging on a flowering plant species; both pollinator and plant were then identified to the species level. Those datasets comprise overall 393 pollinator species, and 392 plant species, with 3890 plant-pollinator interactions (Supplementary Text A1). Prior to calculating diet breadth, we rarefied the data to 20 visitation records per bee species, to avoid confounding rarity with specialization (Blüthgen et al. 2008; Winfree et al. 2014). Nine species had fewer than 20 records and we were unable to estimate diet breadth in the manner described above. Five of these species are known to be specialized and we simulated the diet breadth index of 20 individuals visiting the known host plants. The four other species are known to be generalists and we therefore used the mean diet breadth of its genus. These four species were extremely rare (< 5 records each) in our analyzed dataset.

To calculate diet breadth for each bee species, we considered the number of plants species as well as the phylogenetic breadth that the bees fed upon by using a rarefied phylogenetic diversity index (Nipperess and Matsen 2013). To determine phylogenetic distances among plants, we first constructed a general phylogenetic tree using the PHYLOMATIC “megatree” (version R201120829, Chamberlain and Szöcs 2013) which defines relationships between higher plants (Webb et al. 2008). We then dated nodes across this tree according to Wikström et al. (2001) and used the branch-length adjustment algorithm BLADJ to estimate the age of all remaining, undated nodes. Though this procedure implies that ages within our phylogenies should be treated as approximations (Beaulieu et al. 2007), previous analysis indicates marked improvements of phylogenetic analyses when even a limited number of nodes are properly dated (Webb 2000).

### Statistical analysis

#### Response traits

To investigate which traits are associated with environmental variables related to agricultural intensification, we used a model-based approach to the fourth-corner problem (Brown et al. 2014). The fourth-corner problem highlights the difficulty of studying the environment-trait associations and can be conceptualized as a set of four matrices: abundances by species, trait data by species, environmental data by sites, and environmental data by traits, being the relationships of this last corner the ones to be estimated (Legendre et al. 1997). The core idea of the model-based approach is to fit a predictive model for species abundance as a function of environmental variables, species traits and their interaction. The environment-trait interaction coefficients can be understood as the fourth corner and describes how environmental response across taxa varies as traits vary. The size of coefficients is a measure of importance and are interpreted as the amount by which a unit (1 sd) change in the trait variable changes the slope of the relationship between abundance and a given environmental variable. To estimate these coefficients, we used a LASSO-penalised negative binomial regression (R package “mvabund”, Wang et al. 2012). The LASSO penalty simplifies interpretation because it automatically does model selection by setting to zero any interaction coefficients that do not help reduce BIC. A species effect is included in the model (i.e. a different intercept term for each species), so that traits are used to explain patterns in relative abundance across taxa not patterns in absolute abundance. Pseudo-R2 is calculated as the R2 of the predicted against the observed abundance values for each species at each site.

#### Effect trait analysis

To determine which traits influenced functional contribution of each species, we ran separate linear models with either visitation or per capita efficiency as response variables. Species traits were predictors. The best model based on AICc was selected. When differences between the best models were less than 2 we selected the simpler model.

The analysis for efficiency was done at the species group level (see above: pollination function section). To obtain traits at the species-group level, we calculated the mean values over species belonging to the same group, weighted by the species mean abundance within the group. For categorical variables we chose the dominant level, again weighted by species abundance. This way, we assure that while species within a functional group are selected to be functionally similar, the average traits used reflects species composition.

All residuals were visually inspected to validate model assumptions. All statistical analyses were performed in R (version 3.0.3, <www.r-project.org>).

## Results

### Response traits

Overall, we did not find a strong correlation between any ecological traits and the environmental variables analyzed despite finding a general response of species abundance to change with one or more land use variables (watermelon: estimate of percentage open habitat at 300m = 0.12; blueberry: estimate of percentage agricultural habitat at 300 m= −0.26 and at 1500m = −0.12; cranberry: estimate percentage agricultural percentage habitat at 1500m = −0.23. Supplementary Table A3). Traits do not modify these slopes in most instances, and despite some traits exhibiting weak responses to land use in some cases, these responses were not consistent across crops (Fig 1). For watermelon (overall pseudo-R2 = 0.54), small bees and parasites tended to decline with increasing percentage of agriculture at 300m radius (Interaction estimate of % agriculture at 300m with body size = 0.19, Fig 1D; and with Parasitism = 0.10) and parasites also declined with increasing open areas at 1500m radius (interaction estimate = 0.13). For blueberry (overall pseudo-R2 = 0.22) short-tongued species increased with increasing agriculture at 1500m (interaction estimate = −0.30). In cranberry (overall pseudo-R2 = 0.59), bees nesting in wood and generalist bees tended to increase with increasing open areas at 300 m (interaction estimate = 0.14 and 0.11 respectively) and bees nesting in soil and bigger bees tend to increase with increasing open areas at 1500 m buffer (interaction estimate = 0.14). A complete list of all comparisons is presented in Supplementary material (Table A3).

**Fig. 1:**
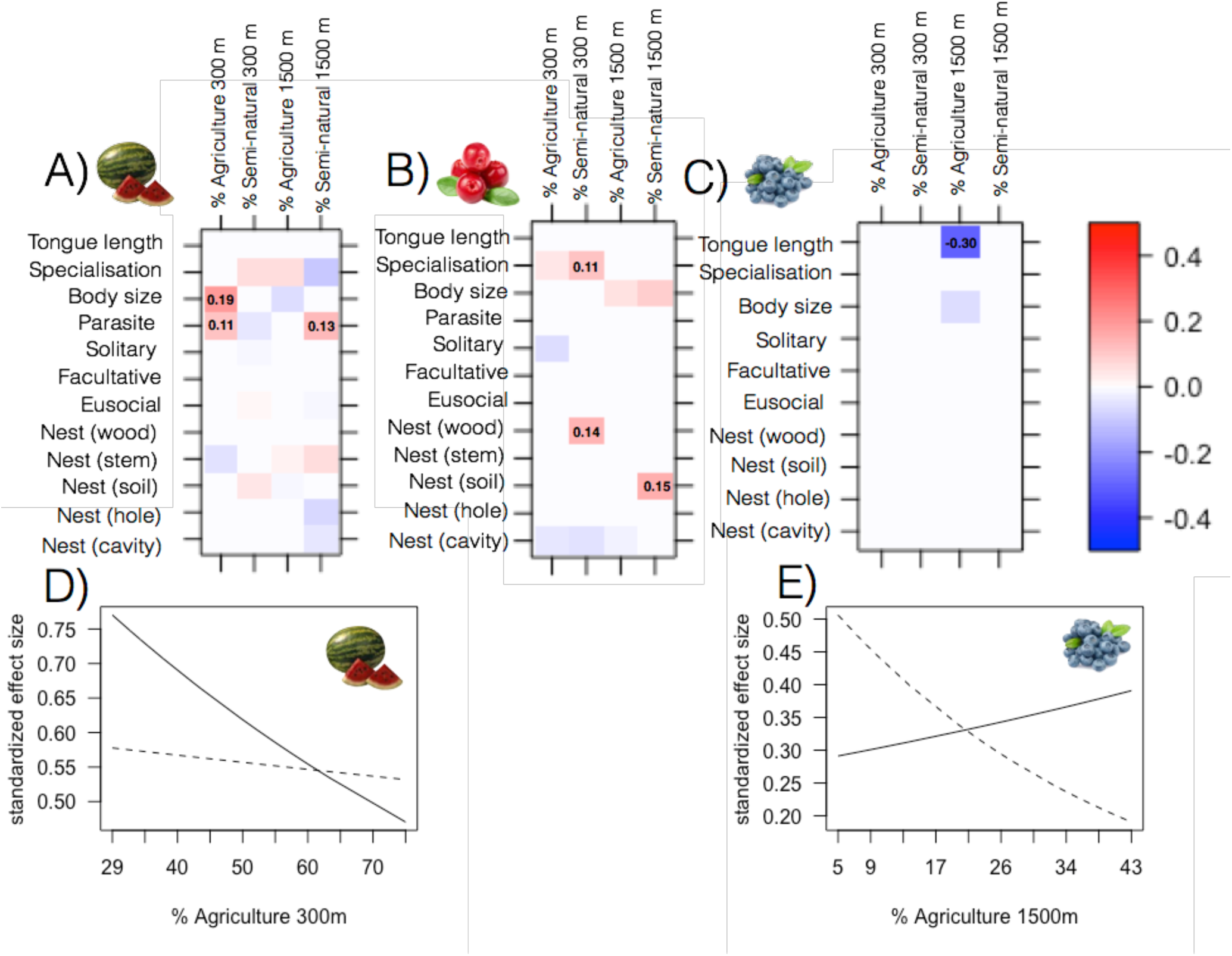
**Relationships between traits and environmental variables** for A) watermelon, B) blueberry and c) cranberry. Positive estimates are in red and negative estimates in blue. Note that the LASSO penalty has set many estimates to zero. D) Detail of the two stronger interactions between body size and percentage of agriculture at 300 meter radii for watermelon and tongue length and % of agriculture at 1500 meter for blueberry. The solid line is the prediction for the 25 percentile of body size and tongue length, while the dashed line is the prediction for the 75% of body size and tongue length for watermelon and blueberry respectively.

### Effect traits

As for response traits, no traits were highly predictive of either visitation frequency or per visit efficiency across crops. For watermelon, the best model for visitation frequency does not includes any trait. However, per visit efficiency was positively correlated with body size and tongue length (R2 = 0.75, F2,9 = 17.07, p < 0.001, Fig 2A). For cranberry, visitation frequency was positively related to cavity nesters (R2 = 0.38, F4,36 = 7.1, p < 0.0001, Fig 2B). This result was driven by *Bombus* species, which are the only cavity nesters in this data set. In cranberry per visit efficiency was not related to any trait. For blueberry, visitation frequency was positively related to diet specialism (R2 = 0.37, F1,20 = 13.5, p = 0.001, Fig 2C), while efficiency per visit is positively related to tongue length (R2 = 0.70, F1,5 = 14.9, p = 0.01, Fig 2D). Model selection, can be found in Supplementary material (Table A4).

**Fig. 2:**
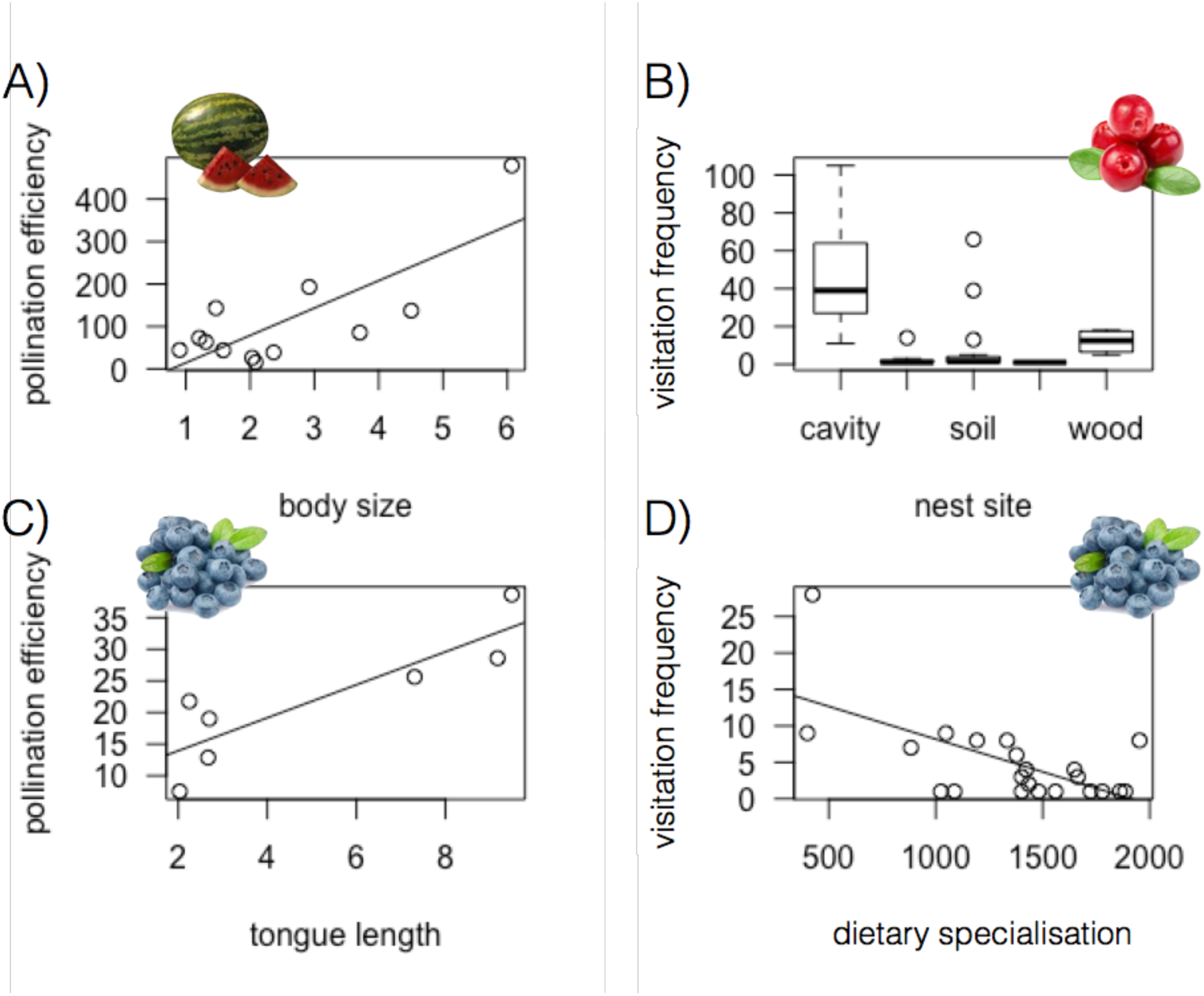
**Multipanel plot showing the relationships between species traits and pollination function**, which is decomposed into efficiency (pollen deposited per flower visit) and frequency of flower visits. A) watermelon, B) cranberry, C-D) blueberry.

## Discussion

Identifying traits that characterize which species are more sensitive to land-use change or those that are functionally important is complex. We found some evidence for response and effect traits but they differed among crop species as well as landscape variable used. Therefore, while some traits may be important in some contexts, no traits were generalizable enough to be used to predict how land-use change will influence the delivery of pollination services across these systems. Further, the relationships identified were weak. This does not negate the importance of traits for understanding which mechanisms underlie species responses to land-use change or pollination effectiveness, but it does suggest that traits commonly used for wild bees might not be suitable for predicting which species will decline or how land-use change will influence the delivery of ecosystem services. In fact, the trait-based literature in general is characterized by weak and/or idiosyncratic relationships between traits and either species responses and functional effects (Tables 1 and 2).

**Table 1:** Summary of some recent studies identifying response traits and its relationship with environmental variables. Environmental variables have been grouped in two main categories because each study uses different metrics. Habitat loss (e.g. isolation, % natural habitat, habitat fragment size) and Agricultural intensification (e.g. natural vs agricultural, % agriculture). Only the direction of the response is indicated, as the different analysis makes any comparison of effect sizes meaningless. Note that in addition, most of this relationships are weakly predictive. See text for details.

|  |  | Relationship direction |  |  |  |
| --- | --- | --- | --- | --- | --- |
| Trait | Environment | Positive | Neutral | Negative | Context dependent |
| Body size | Habitat loss | 0 | 1<sup>9</sup> | 2<sup>1,10</sup> | 1<sup>2</sup> |
|  | Agricultural intensification | 2<sup>7,11</sup> | 2<sup>3,6</sup> | 4<sup>1,4,5,8</sup> | 0 |
| Diet specialization | Habitat loss | 1<sup>3</sup> | 0 | 2<sup>1,9</sup> | 1<sup>2</sup> |
|  | Agricultural intensification | 0 | 1<sup>8,6</sup> | 4<sup>3,7</sup> | 0 |
| Sociality (social) | Habitat loss | 0 | 1<sup>2</sup> | 1<sup>3</sup> | 1<sup>10</sup> |
|  | Agricultural intensification | 4<sup>3,4,8</sup> | 2<sup>6,7</sup> | 0 | 0 |
| Nest location (below-ground) | Habitat loss | 0 | 1<sup>3</sup> | 0 | 0 |
|  | Agricultural intensification | 0 | 2<sup>3,6</sup> | 2<sup>8,11</sup> | 1<sup>7</sup> |

<sup>1</sup>Larsen *et al.* 2005, <sup>2</sup>Bommarco *et al.* 2010, <sup>3</sup>Williams *et al.* 2010, <sup>4</sup>Rader *et al.* 2014,
<sup>5</sup>Benjamin *et al.* 2014, <sup>6</sup>Forrest *et al.* 2015, <sup>7</sup>De Palma *et al.* 2015, <sup>8</sup>Carrié *et al.* 2016, <sup>9</sup>Cane
et al. 2006, <sup>10</sup>Jauker et al. 2013, <sup>11</sup>Klein et al 2008.

**Table 2:** Summary of some recent studies identifying effect traits and its relationship with ecosystem functioning. Only body size is included, as other traits are rarely measured (but see Stavert et al 2016 for hairiness). Only the direction of the response is indicated, as the different analysis makes any comparison of effect sizes meaningless. Note that in addition, most of these relationships are weakly predictive. See text for details.

| Trait | Function | Relationship direction |  |  |  |
| --- | --- | --- | --- | --- | --- |
|  |  | Positive | Neutral | Negative | Context dependent |
| Body size | Pollen deposition | 2<sup>1,2</sup> | 1<sup>3</sup> | 0 | 0 |
|  | Visitation rate | 0 | 0 | 1<sup>2</sup> | 0 |
|  | Fruit set | 0 | 0 | 0 | 2<sup>4,5</sup> |

<sup>1</sup>Larsen *et al.* 2010, <sup>2</sup>Hohen et al. 2008, <sup>3</sup>Stavert et al. 2016, <sup>4</sup>Garibaldi et al. 2015, <sup>5</sup>Gagic et al. 2015

Being able to identify strong response traits would be a key tool for understanding extinction risk, and an asset for conservation planning. However, characterizing extinction risk based on traits is challenging. Despite some generalities that emerge across taxa, with rare species, big species, specialists, and higher trophic levels being in general more sensitive to disturbances (Fisher and Owens 2004), there is a large variation in the response of the species with those traits (Fritz et al. 2009; Séguin et al. 2014). Work specifically on native bees has found that traits such as specialization, body size, and sociality may predict responses to land use (Table 1; Winfree et al. 2009, Bommarco et al. 2010, Williams et al. 2010, Bartomeus *et al.* 2013b, Kremen and M’Gonigle 2015, De Palma et al. 2015, Carrié et al. 2016). However, studies often find contrasting results (Table 1). For example, De Palma et al. (2015) analyzed over 70,000 wild bee records and found that small species were most sensitive to agricultural land use, while others have found that larger species are more sensitive to agricultural land use and/or environmental change generally (Larsen et al. 2005; Bartomeus et al. 2013b), and some have found little effect of body size (Williams et al. 2010). Here, we found a weak trend for small species to be more sensitive to local agricultural intensification in watermelon, but this trend disappears when land use is measured at larger scales. Another trait, dietary specialization, is one of the few traits that has been generally linked to increased species sensitivity to environmental change (Table 1, Williams et al. 2010; Scheper et al. 2014; De Palma et al. 2015), but here we found that floral specialist bees did not decline with intensifying agriculture. If anything, one of the most abundant bee species in the cranberry system (*Mellita americana*) is a specialist on cranberry (*Vaccinium macrocarpon*). Specialist bees observed in crop systems are likely to be specialized on the crop plant family as was the case in our data (e.g. *Mellita americana* in cranberry, but also *Habropoda sp.* and *Andrena bradleyi* in blueberry and *Peponapis pruinosa* in watermelon). We would expect different responses from study designs that include natural habitat and a larger range of specialist host plants (Forrest et al. 2015, Bartomeus and Winfree 2013).

Alternatively, the lack of strong trait-environment associations may be due to the variables used to measure agricultural intensification being too coarse to detect common responses. While finer-resolution studies will undoubtedly be informative, they are unlikely to lead to a greater likelihood of predicting how changes in biodiversity affects the delivery of ecosystem services if these measures are difficult to quantify or are context dependent.

Effect traits have been even harder to identify for pollinators. The limited data published on particular plants suggests insects with larger bodies tend to deposit more pollen per flower visit, but this pollen was not well distributed on the stigma (Table 2; Hoehn et al. 2008), or that the correlation between body size and per visit pollination function is low (Larsen et al. 2005). Our study supports the positive correlation between body size and per-visit pollen deposition in both watermelon and blueberry (although note that tongue length is correlated with body size in blueberry r = 0.76), but not for cranberry. Hence, generality is difficult to achieve because a single pollinator trait, like big body size, may not lead to high pollination function in all contexts. Rather it seems likely that the most efficient trait will depend on the crop (Garibaldi et al. 2015). Moreover, the total pollination provided by a pollinator species is the product of visitation frequency and per capita efficiency (Kremen et al. 2005), two processes that may be governed by different traits.

If generalizable response and effect traits can be found, the final step will be to link response and effects to predict changes in ecosystem services. A positive association between the response and effect traits (Naeem and Wright 2003) such that species with the strongest response to environmental change also had the strongest effect on function, indicates the land-use change has the potential for dramatic effects on ecosystem function. Whether response and effect traits are in general positively, negatively, or uncorrelated is an important question that has not yet been answered (Larsen et al. 2005). Despite the conceptual elegance of the response-effect trait framework, it is only effective if it is predictive, and strong evidence for the generality of traits has not yet been found. For example, even the very thorough and rigorously analyzed study of response-effect relationships by Larsen et al. (2005) is based on a non-significant weak relationship between pollinator per visit efficiency and body size. Similarly, the marginal R^2^ (i.e. variance explained by fixed effects) of the best model including traits in the comprehensive analysis done by De Palma et al. (2015) is lower than 0.1. Similarly, in our study, even the strongest correlations found for watermelon, where big species are less sensitive to local agricultural intensification and more efficient per visit, but not more frequent flower visitors than smaller species are too weak to be useful for predictive purposes.

Predictive response and/or effect traits are often assumed in the larger literature as well. For example, recent re-evaluations of community stability in food webs shows that using body size as proxy of extinction risk changes the outcome of the stability simulations (Brose et al. 2016). However, the assumption that body size is a good predictor of extinction risk is not directly validated. Given the correlation showing that bigger species are more sensitive is usually weak (Fisher and Owens 2004), these kind of approaches could produce misleading outcomes.

Currently trait data may be too coarse to reveal ubiquitous response and effect traits for four reasons. First, some traits may simply reflect identity of genera or higher taxonomic groups. For example, some bumble bee species in our three systems (especially *B. impatiens*) are common, functionally dominant, and robust to extinction (Cariveau et al. 2013, Winfree et al. 2015). Some of the important response and effect traits that we found, such as cavity nesting and body size, may simply be proxies for bumble bees. Bumble bee species also share other traits (e.g. sociality) that are commonly used in trait analyses. Therefore, studies that don’t include phylogenetic correlations may be simply characterizing the general relationship between disturbance and the functionally dominant taxa. As there is a great variability in the responses to disturbance among bumble bee species (Cameron et al. 2011; Bartomeus et al. 2013b, Persson et al. 2015) this may also explain why some studies find big species to be more sensitive to land-use change (Larsen et al. 2005) and other studies find the opposite (Rader et al. 2014, this study for watermelon). Second, traits may interact in complex ways and single traits may be not able to capture responses and functional contributions across species (e.g. Bommarco et al. 2010). Third, phenotypic variability within species, usually ignored in trait-based approaches, may play a more important role than previously though (Bolnik et al. 2011). Finally, the most important traits may not have been studied. Response traits such as dispersal ability, fecundity, and nest microclimate/soil type, and effect traits like floral visitation behavior or hairiness (Stavert et al. 2016) may be better predictors than the traits we have now. However, if these traits are not easy to measure across bee species, they may be of little use. Traits databases that include an increasing number of traits and agreed-upon measurement techniques similar to those used in plant ecology (Kattge et al. 2011) but that are also open-access may lead to significant advancements in functional trait ecology in wild bees.

There is a call to be more predictive in ecology (Petchey et al. 2015, Houlahan et al. 2017). The use of traits to predict species responses and subsequent changes in ecosystem services is a potentially powerful approach. This is especially the case for organisms such as insects where species identification is challenging and detailed species-level natural history information is lacking. The ability to effectively use a trait framework is becoming controversial because studies thus far have not clearly related specific traits to specific threats or functions (Didham et al. 2016; Shipley et al. 2016). A growing number of studies are working to address the complexity and increase the predictability of this framework (e.g. Laughlin and Messier 2015). However, until these approaches yield consistent patterns across systems, site-specific species identity and monitoring may at present be the best measure for predicting changes in ecosystem services as a result of land-use change. A few dominant species often drive ecosystem functioning (Kleijn et al. 2015; Winfree et al. 2015). Identifying the sensitivity of the functionally dominant species may be the best proxy thus far for predicting effects of species loss in ecosystem function.

## Acknowledgments

We thank Scott Chamberlain for help with BLADJ and Paco Rodriguez for statistical advice. IB was supported by project Beefun (PCIG14-GA-2013-631653) funded by EU.

## Data Accessibility

All data and code used in this manuscript is accessible in github (https://github.com/ibartomeus/RE_traits) and will be archived on acceptance in figshare.

## Supplementary material

### Text A1

Datasets used for calculating dietary specialization: Six datasets were used to create the phylogenetic distance index. All data were collected in the region of the crop study. Specimens were collected using a hand net and the bee species and plant species were recorded. This resulted in a total of 18,733 bee x plant interactions for species that were also in the crop dataset. The number of species, sites, and years of collection are as follows: 1) Pine barrens in 2003: 280 bee x plant interactions. Habitat types were extensive pine-oak forest (14 sites), forest fragments (14 sites), suburban back yards (7 sites), and agricultural field borders (5 sites) in New Jersey (Winfree et al. 2007). Bees were collected in temporally stratified sampling rounds between April and September. 2) NJPA: 3906 bee x plant interactions. Data collected on watermelon field margins at a total of 20 sites. Farm types included small-scale mixed farming, both crops and field margins, both organic and low-pesticide-input conventional. All bees were collected in three temporally stratified sampling rounds in July, in each of 3 years. 3) NFWF 3906 bee x plant interactions. Habitat types were old fields. Bees were collected in May through Sept at 25 sites for two years. *Lasioglossum* species where not included for this dataset due to recent changes in its taxonomy. 4) NSF 2006 666 bee x plant interactions. Habitat types were deciduous forest fragments (13 sites), and suburban / urban yards (3 sites) and sites with extensive forests with diverse wildflower communities (4 sites). All bees were collected in sampling rounds between April and early June. 5) CIG 4600 bee x plant interactions. Site were comprised of old fields as well as pollinator enhancement sites. Bees using were collected using a hand net from a total of a total 18 sites in 2011-2013. For each bee specimen, the plant species was recorded. 6) Cape May 5858 bee x plant interactions. This study included only one site. The habitat was an old field that had been planted in 20 species of native perennial plants. Sampling took place over 3 years in sampling rounds that occurred in May through September.

Winfree, R. Griswold, T. and Kremen, C. (2007). Effect of human disturbance on bee communities in a forested ecosystem. *Conservation Biology*. 21: 213–223.

**Table A1:**
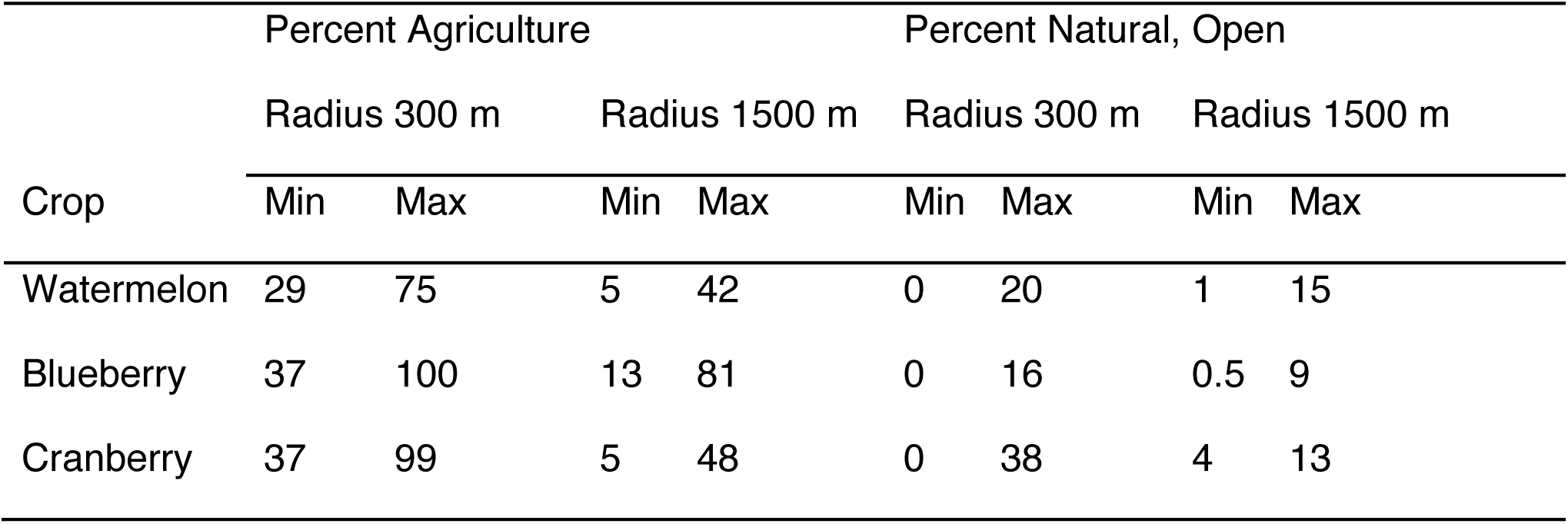
Range of variation in agricultural and semi-natural land cover for three crop systems

| Crop | Percent Agriculture |  |  |  | Percent Natural, Open |  |  |  |
| --- | --- | --- | --- | --- | --- | --- | --- | --- |
|  | Radius 300 m |  | Radius 1500 m |  | Radius 300 m |  | Radius 1500 m |  |
|  | Min | Max | Min | Max | Min | Max | Min | Max |
| Watermelon | 29 | 75 | 5 | 42 | 0 | 20 | 1 | 15 |
| Blueberry | 37 | 100 | 13 | 81 | 0 | 16 | 0.5 | 9 |
| Cranberry | 37 | 99 | 5 | 48 | 0 | 38 | 4 | 13 |

**Table A2:**
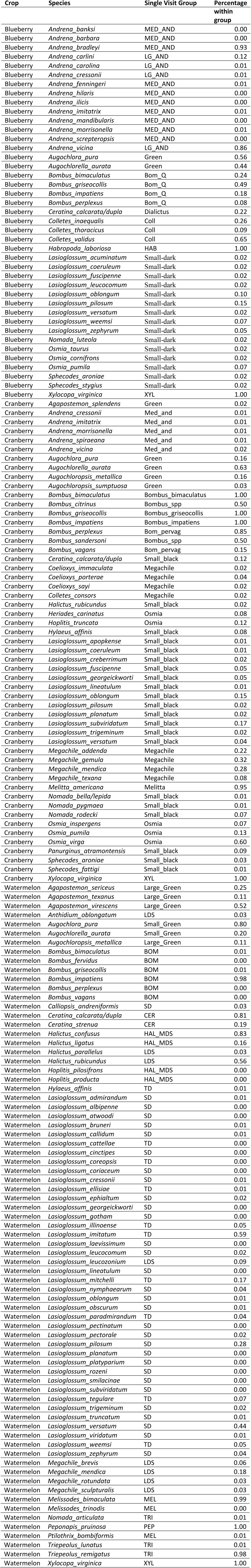
Equivalencies between species and groups used for single visit data.

**Table A3:**
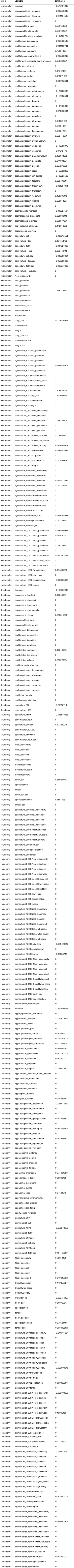
Response trait model estimates for all variables, including the fourth corner interactions. Note that many coefficients are set to zero due to the lasso penalty which acts as model selection.

**Table A4:** Model selection procedure showing all models within 2 AICc values.

| Crop | Measure | Model | Delta<br>AICc |
| --- | --- | --- | --- |
| Watermelon | Visitation Frequency | ~ body size + tongue | 0 |
| Watermelon | Visitation Frequency | ~ tongue | 0.06 |
| Watermelon | Visitation Frequency | ~ . | 0.28 |
| Watermelon | Visitation Frequency | ~ body size | 1.99 |
| Watermelon | Pollen deposition | ~ body size + tongue | 0 |
| Cranberry | Visitation Frequency | ~ sociality + tongue | 0 |
| Cranberry | Visitation Frequency | ~ nest place | 1.43 |
| Cranberry | Visitation Frequency | ~ sociality | 1.56 |
| Cranberry | Pollen deposition | ~ . | 0 |
| Blueberry | Visitation Frequency | ~ specialization | 0 |
| Blueberry | Pollen deposition | ~ tongue | 0 |

